# Vascularized Brain Organoid: A Versatile Platform Models Brain Cancer and Traumatic Brain Injury

**DOI:** 10.64898/2026.08.11.744207

**Authors:** Shu-Wei Angela Huang, Chin-Hsing Annie Lin

**Author notes:** Corresponding author: Chin-Hsing Annie Lin, Ph.D. University of Texas at San Antonio One UTSA Circle San Antonio, TX78249.

## Abstract

Human iPSC-derived brain organoids are revolutionizing tools to study layers biology, synergize disease modeling, and accelerate therapeutic discoveries that overcome obstacles in monolayer cell culture or animal models. The neurovascular unit including vasculature and microglia is critical for brain development, maintenance of synaptic plasticity and neural activity, and the high metabolic demands of long-term culture. We present a methodology to incorporate these important components during organoid generation and discuss potential approach, aiming consistent production of vascularized organoids for longitudinal study. We also demonstrate that this vascularized organoid is a versatile platform to model brain cancer and traumatic brain injury.

## INTRODUCTION

The breakthrough with human iPSC in culture enables 3D self-organization processes inherent to organogenesis, giving rise to biomimetic systems termed ‘‘organoids’’. Leveraging developmental nature of stem cell hierarchy, different types of organoids were successfully generated to represent a plethora of organs and tissues including brain organoids ^1–4^. These organoids are new frontiers in understanding human brain development, modeling neuropathological disorders, and developing therapeutics. While organoids are entering their heyday, challenges remain regarding these models fail to capture microenvironment and physiology, critical for sustaining neural activity and long-term survival. Absence of neurovascular unit (vasculature and microglia) in brain organoids is a hurdle poised to transform both basic research and translational medicine. Vasculature is vital support for the high metabolic demands of continuous neurogenesis and synaptic activity, while microglia regulate neuron survival and synapse pruning. Hence, vascularization and incorporation of microglia facilitate functional integration of new-born neurons in brain organoids to sustain spontaneous activity and respond to external stimulation. In this work, a well-integrated protocol capitalizes on developmental principle and bioengineering approach to generate vascularized brain organoid with microglia as the sophisticated modeling of brain cancer and traumatic brain injury (TBI). The architecture in the model will provide opportunities to study cancer neuroscience, assess targeted therapy for brain cancers and TBI, as well as understand pathological condensates and senescence in disease or injury conditions.

## MATERIALS and METHODS

### Human iPSC culture

The iPSC lines (Stem Cell Technology) were generated from mononuclear cells (MNCs) isolated from healthy donors’ peripheral blood cells. The quality of iPSCs were examined based on morphology, genomic integrity, pluripotency tests, and marker gene expression that was accomplished using the hiPSC Genetic Analysis Kit, karyotyping (KaryoStat Assays; Thermo Fisher), STEMdiff Tri-lineage Differentiation Kit, and Tri-lineage Differentiation qPCR Array, respectively. The iPSC culture was performed by seeding human iPSC (hiPSC) 1×10^5^ to 12-well plate or 0.5×10^5^ ∼1×10^6^ to 6-well plate using mTeSR1 medium (STEMCELL Technologies) containing ROCK inhibitor 10 μM Y27632 (STEMCELL Technologies) and cultured in either feeder-dependent or feeder-independent plates. Alternatively, iPSC cells were cultured in human ESC medium (DMEM/F12), 1% Glutamax (Invitrogen), 20% [v/v] Knockout Serum Replacement KOSR (Invitrogen), 1% [v/v] MEM-NEAA (Invitrogen), 100μM 2β-mercaptoethanol (Invitrogen), 15% Fetal Bovine Serum (Invitrogen),10 μM Y-27632 and 4ng/mL thermostable human basic fibroblast growth factor (bFGF, Invitrogen).

### Feeder-dependent

Plates were coated with Matrigel for 1 hour at room temp (Corning or Bioscience: growth-factor depleted Matrigel, 1:125 diluted in DMEMF/F12 medium, equivalent to 80mg/ml Matrigel), aspirate extra Matrigel before seeding iPSC. Note: Each lot of Matrigel has its certification. The endotoxin level should be lower than 1.5 unit in certification.

### Feeder-independent

Plates were coated with 0.1% (w/v) gelatin for at least 20 min (autoclaved to dissolve gelatin and keep at sterile condition).

Human iPSCs were maintained in mTeSR1 medium or human ESC medium with ROCK inhibitor and passaged using Accutase (STEMCELL Technologies). After ∼70% confluence, hiPSC colonies were dissociated into single cells with 1ml Accutase for 3∼4 minutes incubation at 37°C to make sure single cell suspension. Then, adding medium to dilute Accutase and using centrifuge to gently spin down iPSCs. Cell count was performed by mixing 2uL cells and 18uL trypan blue and loading 10uL onto hemocytometer (Adjust dilution if hemocytometer count is too low/high).

*Freeze early passage of iPSC lines:* mycoplasma testing was performed prior to freezing iPSCs using iPSC medium plus 5%DMSO or FreSR^TM^-S cryopreservation from Stem Cell Technologies (0.5×10^5^ ∼1×10^6^ cells/vial; passage 1).

### Generation of human endothelial cells (HEC)

iPSC colonies were dissociated with Accutase. One million of cells were cultured in 5mL STEMdiff APEL2 medium (Stem Cell Technologies) in the presence of 10μM Y-27632 (Stem Cell Technologies), 6μM CHIR99021 (BioTechne), and recombinant proteins 50ng/mL human VEGF and 50ng/mL BMP-4 (PeproTech) in a 25 cm^2^ flask for 4-5 days.

### EB formation and vascularized brain organoids generation

To generate vascularized brain organoid, first, iPSCs were seeded at a maximum density of 25 cells/cm^2^ and were maintained in feeder-independent human stem cell culture medium containing bFGF and ROCK inhibitors until embryoid body (EB) formation prior to induction of neural ectoderm.

Approximately 7000∼9000 hiPSCs were resuspended in fresh-made mTeSR1 medium or human ESC medium containing ROCK inhibitor 10 μM Y27632, and then seeded into the low-attachment U-bottom 96-well plate (Thermo), 150 μl per well to form aggregate EBs. When the size of EB reached ∼400μm, EB were cultured in medium without bFGF or Y-27632 for 6 h. Then, a single EB was co-cultured with dissociated HEC for 2 days in neural induction medium (DMEM/F12, 1% Glutamax containing 1% [v/v] N2 (Invitrogen), 1% [v/v] MEM-NEAA, 1μg/mL Heparin (Stem Cell Technologies). During neural induction toward neural ectoderm, we introduced transcription factor PU.1 recombinant protein to generate microglia-like cells in brain organoids^5^. On day 2, the co-culture of EB-HEC complex was embedded into 30μL droplet of fibrin gel ^6^ and cultured with cerebral organoid differentiation medium (50% [v/v] DMEM/F12, 50% [v/v] Neurobasal (Invitrogen), 1% [v/v] Glutamax, 0.5% [v/v] N2, 2.63 μg/mL Insulin, 0.5% [v/v] MEM-NEAA, 50μM 2β-mercaptoethanol, 0.5% [v/v] B27 without vitamin A (Invitrogen), 20ng/mL VEGF-A, 100ng/mL IL-34 (PeproTech), 10ng/mL GM-CSF (PeproTech). After 2–3 days, the co-culture medium was replaced by cerebral organoid differentiation medium with 0.5% (v/v) B27 containing vitamin A (Invitrogen), 20ng/mL VEGF-A, 100ng/mL IL-34, 10ng/mL GM-CSF for 3-4 days. The co-cultures were kept under orbital shaker (85 rpm) in the 5% CO2 incubator at 37°C for expansion. After the first media exchange, vascularized brain organoids were continuously maintained in differentiation medium containing vitamin A and VEGF, which was replenished every 7 days to support their growth and expansion. To sustain multilayer vessels and sprouting, we utilized syringe pump driven perfusion (5μL/min) of endothelial cells around growing organoids culture. When necessary, we will add Aprotinin (10 μg/mL) to prevent fibrin degradation on the second week of vascularization process.

### Immunohistochemistry and whole mount imaging

Immunohistochemistry for cryo-sections: vascularized brain organoid and biomimetics was fixed with 4% paraformaldehyde/PBS, cryo-protected with 30% sucrose/PBS prior to cryo-sectioning and immunohistochemistry. The z-stack images were acquired under Zeiss or Nikon confocal microscope and quantified by ImageJ. Whole-mount imaging: we utilize a combination of tissue clearing with Single Molecule Localization Microscopy and light-sheet microscopy to visualize cells within thick and intact tissues in 3D organoid context. A modified iDISCO Tissue Clearing protocol ^7^ was used to eliminate lipids and molecules that renders tissue transparence, allowing light to penetrate into sample without scattering. Using a thin sheet of light to illuminate a single plane within a 3D sample, light-sheet microscopy can minimize photobleaching for live vs prolonged high-speed meso-scale imaging within large tissues. Briefly, brain organoids were fixed with 4% paraformaldehyde/PBS prior to tissue clearing via methanol dehydration series and overnight 66%:33%% methanol: dichloromethane (DCM) incubation, and then 100% DCM and dibenzyl ether (DBE). Cleared tissues were imaged on the UltraMicroscope II light sheet microscope (LaVision, Bielefeld, Germany) and processed with Imaris (Bitplane) software. All steps of imaging quantification were performed in ImageJ following manufacturer’s instruction.

### Microfluidics-based single cell RNA-Seq delineates major cell type in vascularized brain organoids

We previously reported using Printed Droplet Microfluidics (PDM) platform for single nuclei RNA-Seq to assess differential gene expression in dividing cells isolated from hippocampus of postnatal brains ^8^. Here, we utilized PDM devices for high-throughput single-cell RNA-Seq to identify major cell types in brain organoids. Microfluidic devices consist of channels to control single cell loading, where the reaction volumes are reduced to the sub-nanoliter scale ^9^. These reduced volumes allow for the efficient partitioning of individual cell for single-cell analysis by using Drop-Seq technology ^10^, which utilizes oligo-capture beads to uniquely bar-code mRNA from each cell prior to next generation sequencing (NGS). Sequencing data obtained from Drop-Seq were processed with uniform manifold approximation projection (UMAP) to obtain cell-lineage clusters from the samples ^11,12^. Clusters were further annotated as specific cell-types based on expression analysis ^13^.

### Modeling traumatic brain injury via mechanical injury on vascularized brain organoids

We utilized ultrasound within a range of intensity (0.1, 0.6, 1,1.2 MPa) to inflict mechanical injury on vascularized brain organoid, modeling moderate sport-related injury at 0.1MPa and mimicking bomb shockwave ≥ 0.6 MPa. The biomimetics received 1.2 MPa were deformed; thus, experimental design was focus on 0.1, 0.6, and 1MPa.

### Generation of patient-derived GBM biomimetics as pre-clinical and personalized models

De-identified human GBM tumors from surgical resection were dissociated into single cell suspension. Subsequently, 10-40 tumor cells were embedded to the vascularized brain organoid in a single droplet of 1% Matrigel (∼40 μL) as close as possible for fusion process. All failed fusions were removed by aspiration while successfully fused GBM tumor organoid were grown in 10 cm culture dishes containing differentiation medium with 1% Matrigel to facilitate their growth and expansion towards a fully formed GBM biomimetics.

### Validating patient-derived GBM Biomimetics

Molecular profiling is an optimal strategy to compare patient’s tumor with patient-derived translational model. To validate GBM biomimetics recaptures molecular signatures in patient’s tumor, cells isolated from patient’s tumor and derivatives-tumor portion of GBM biomimetics were subjected to RNA-Seq and Ribo-Seq analysis to compare gene expression pattern. Paired-end RNA-Seq 76 base pair reads were aligned to hg38. Count matrices are compatible with widely-used open-source analysis software packages. After normalization, FPKM (fragments per kilobase of exon per million reads mapped) were subjected to unsupervised heatmap comparison between primary tumor and its derived GBM biomimetics.

## RESULTS

The lack of vascularization in human organoids is one of limitations that hamper the use of this *in vitro* model for the study of neuropathology and disease modeling. Based on developmental principle, mesoderm-derived angioblasts give rise to endothelial cells, which migrate into neuroepithelium regions to form primitive blood vessels and then grow into vascular networks. Using this developmental insight as leverage, we modified existing brain organoid culture protocols^1–4^ to promote the integration of immune cells, vessel and brain organoids that enhance organoid expansion and survival. We validated the process of vascularized brain organoid cultures with neuronal and vascular markers by immunohistochemistry (Figure 1). A successfully formed 25-day brain organoid can reach up to the size of 4-5mm (Figure 2A). Immunostaining of cryosections obtained from 25-day brain organoids reveals an increase in mature neurons (NeuN^+^) and interneurons (Calbindin^+^) (Figure 2B), along with CD31^+^ vascularization (Figure 2C). In addition to immunohistochemistry, we utilized transcriptional data as a metric. The mapping of cell types presents the reference of cell clusters in normal condition of vascularized brain organoid (Figure 2D) that is the foundation to compare disease conditions in brain cancer or TBI.

**Figure 1.**
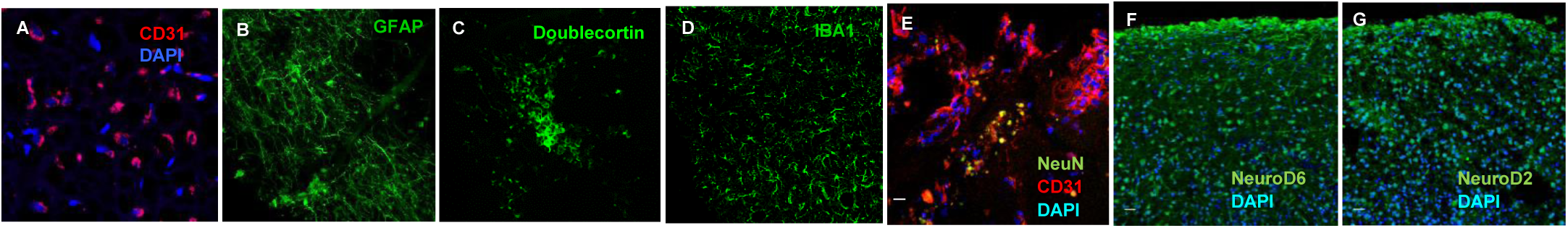
Representative images from the process of vascularized brain organoids. (A, E-G) Immunostaining on cryo-sections and images were acquired by Zeiss 710 confocal microscope. Scale bars= 20μm. (B-D) Whole-mount brain cleaning, immunohistochemistry and imaging. (A) Immunostaining on cryo-section with CD31 antibody showed CD31^+^ endothelial cells and small blood vessels on day-12. (B) GFAP^+^ radial glial cells and (C) Doublecortin ^+^ neuroblasts were present on day-12. (D) IBA1^+^ microglia-like cells were present on day-14. (E) Images from day-15 organoid formation in vitamin A differentiation medium showed CD31^+^ vessel branching, where NeuN ^+^ mature neurons started forming (yellow green signals). On day-20 organoid culture, cortical neuron marker NeuroD6 (F), and corticogenesis and pallial neuron marker NeuroD2 (G) were abundant, demonstrating successful forebrain formation.

**Figure 2.**
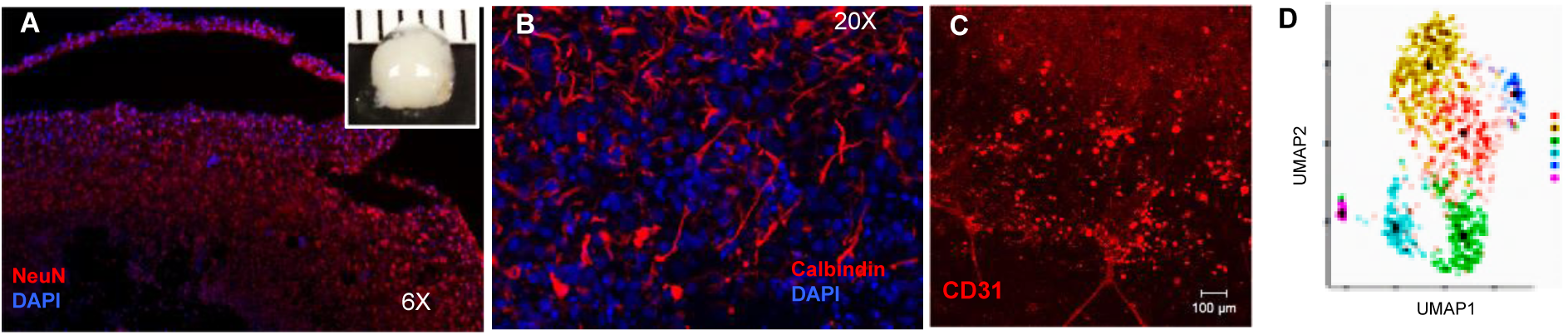
Representative images from day-25 vascularized brain organoid. (A) a culture of 25 days brain organoid reached up to 4∼5mm (inset). Each scale is 1mm. Images taken at 6X showed NeuN^+^ mature neurons were abundant and wildly distributed in 25-day brain organoid. (B) Calbindin^+^ interneurons were abundant in 25-day cryo-section/immunostaining brain organoid. (C) CD31^+^ vasculature on whole-mount imaging demonstrated successful vascularization. Scar bar= 100 μm. (D) Single cell RNA-Seq analysis illustrated at least 6 different cells types in brain organoids.

### Mechanical Injury on vascularized brain organoid showed TBI-related clinical biomarker

To validate the characteristics and reliability of our TBI model, the biomimetics received ≥ 0.6 MPa were subjected to biomarker assessment 4 days post-injury (4dpi). Based on the known patients’ TBI signatures, the hallmark traits of TBI resembling other neurodegenerative diseases that consist multiple phenotypic alterations, such as increases of cell death and phosphorylated TDP43 ^14–22^. We adapted previously described TBI biomarkers and found the accumulation of phosphorylated TDP-43 in cytoplasm of neurons 4 days post-injury (4dpi) at 0.6 MPa (Figure 3).

**Figure 3.**
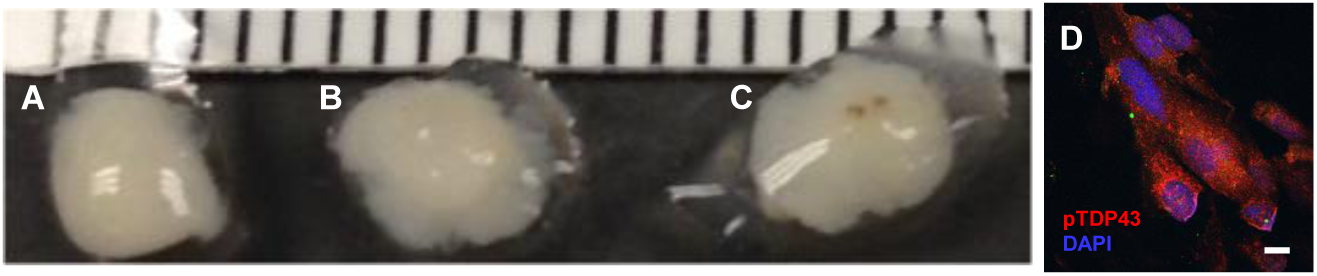
Representative images of TBI model after ultrasound inflicted mechanical injury. (A) 4 days post-injury (dpi4) of biomimetics received 0.1 MPa ultrasound treatment did not show apparent changes; (B) biomimetics treated with 0.6MPa on dpi4; (C) biomimetics treated with 1MPa on dpi4; (D) immunostaining with phosphorylated TDP-43 for biomimetics treated with 1MPa on dpi4 showed accumulation of phosphorylated TDP-43 in cytoplasm, Scale bars= 20μm.

### Cellular and molecular characterization of GBM biomimetics

Overcoming simple GBM organoids composed of only GBM cells, we incorporated GBM tumor cells into vascularized brain organoid (GBM biomimetics) that better mimic tumor microenvironment and interaction among neuron-vessel-tumor. Representative images showed GBM biomimetics; tumor growth and vessel branching were evident (Figure 4A, 4B). Using immunostaining on cryo-sections of GBM biomimetics, CD31^+^ vessel sprouting and less NeuN^+^ cells were found in GBM biomimetics (Figure 4C). In addition, RNA-Seq (Figure 4D) and Ribo-Seq (Figure 4E) analyses illustrated a similar trend of gene expression pattern between primary tumor and its derived GBM biomimetics. The high similarities include genes involved in cell cycle, cell death, and ECM remodeling with equivalent log10-fpkm values (Figure 4D). Although marginal discrepancy was found for some genes in ribosomal biogenesis (log10 value at 5 versus log10 value at 12), the overall trend of increased expression was consistent.

**Figure 4.**
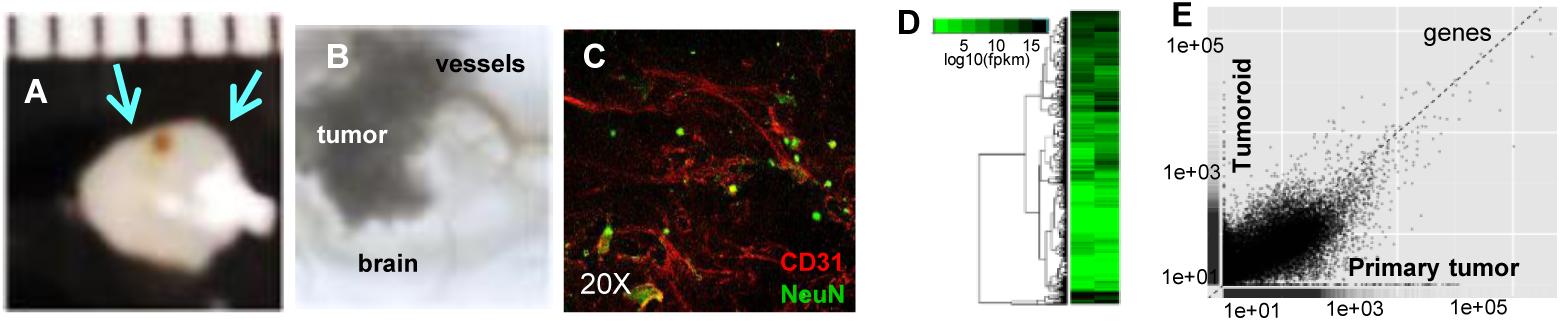
Representative images and molecular characterization of GBM biomimetics. (A) Fusion culture of GBM biomimetics at 45 days. (B) zoom-in image of A showed fusion of tumor in brain organoid with vessels branching. Arrowheads point the area of zoom in. (C) CD31^+^ vessel sprouting is evident and NeuN^+^ mature neurons were less in tumor portion. Arrowheads point the area where image was acquired under Zeiss 710 confocal microscope (20X). (D) RNA-Seq analysis: Unsupervised heatmap illustrated similar gene expression profile between primary tumor and GBM biomimetics. Genes with log 10 fpkm>1 were presented on heatmap. (E) Scatterplot showed positive linear correlation by Ribo-Seq analysis.

## DISCUSSION

Our laboratory-based protocol emphasizes the incorporation of vascular and microglial components during organoid generation rather than adding them as secondary supports. This approach is based on the rationales that the interaction among neuron-vasculature-microglia microenvironment can sustain synaptic metabolism over time. As a result, a well-integrated microenvironment capitalizes on co-existing of neurons, vasculature and glial cells to maximize spontaneous activity. For long-term culture, vasculature improves nutrient delivery into the bulk of brain organoid and reducing the loss of function that often arises when larger constructs become diffusion-limited. Thus, spontaneous neuronal activity is strongest and most durable when brain organoids are built with these supporting features normally present in living condition.

### Neuroscience

We demonstrated the improved architecture of brain organoids with vasculature and glial cells. Relevant to neurobiology, growing evidence supports that biomolecular condensates formed by phase separation play key roles in normal and aberrant conditions, including synaptic density critical for neural communication. This 3D human-based model can be built to understand the mechanism of condensate formation; also, can be designed to perform spontaneous activity without external stimulation and to strengthen electrically evoked spiking with the supported architecture to maximize circuit connectivity and neuronal excitability.

### Disease modeling

Beyond the compatibility of neuroscience research, a sophisticated patient-derived brain tumor organoid model has potential to enhance synergistic research in tumor immunobiology, tumor-neuron interactions, and a robust pipeline of therapeutic drug screening. Our GBM biomimetics displays the primitive microenvironment that represents dynamic cellular and physiological interactions. This model can be maintained for long time in culture without passages that will allow our investigation of neuron-immune-tumor crosstalk and explore multiple therapeutics along with monitoring the full extent of tumor-specific therapeutic efficacy and prediction of recurrence or resistance to the therapy. Neither tumor spheres nor cell culture systems can mimic such cellular diversity and microenvironmental gradients pairing with drug screening. Furthermore, our scalable feature of TBI modeling enables a range of mechanical injury via high-intensity focused ultrasound on a standard multiple-well culture plate. This way can reduce unnecessary variation between pre-clinical tests, making it well suited for high-throughput screening and molecular analyses of TBI, amenable to longitudinal assays across experiments. Lastly, the 3D architecture enables modeling environmental exposure at various doses and time points that relieve intensive labor and animal use.

### Bioengineering in building organoid

When tissues are generated by uncontrolled aggregation, differences in performance can reflect variability in the starting material rather than meaningful differences in architecture, behavior, network composition, or neurocircuit vs external response. Consistency and sustainability of organoid production are essential to addresses a central challenge in organoid-based modeling or computational biology. The single-cell deterministic printing and microfluidics technologies can complement our improved protocol. Printing provides precise control over starting composition and structure that preserve consistent organoids, while microfluidic culture provides continuous exchange of nutrients and metabolites (i.e., amino acids of neurotransmitter precursors), creating a more stable environment for sustained neuronal signaling. This is especially important for maintaining spontaneous activity, because neurons require access to metabolic substrates and benefit from controlled removal of waste products that would otherwise degrade tissue performance over time. In addition to fibrin gel, lithographical fabrication of collagen hydrogels (7.5mg/ml) can optimize vessel formation to sustain organoid culture. Where feasible, the microfluidic configuration can also be extended toward an air-liquid interface format to improve oxygen support and further reduce necrotic effects in long-term culture. Together, combining single-cell printing and microfluidics will create a practical route to producing reproducible brain organoids that are not only alive, but capable of sustained spontaneous activity and reliable response to stimulation. Future work will leverage these established methods and guided patterning strategy to assembly distinct regions of brain organoids with functional properties tailored for biological and biomedical research.

## CONCLUSION

Future direction to refine the approach will combine controlled assembly with guided developmental patterning by using region-specific signaling molecules and transcriptional factors to direct formation of defined neuronal identities, rather than relying solely on self-aggregation. This provides a path to generating related but distinct organoid architectures whose maturation state, cellular composition, and circuit properties can be tuned with different functional objectives.

## ACKNOWLEDGMENT

We thank Dr. Adam Abate for insightful discussion of bioengineering approach. Authors are grateful to Drs. Eric Brey and Feipeng Yang for technical suggestion as well as Drs. Chris Rhodes and Chenwei Lin for data analysis.

## Author contributions

SAH performed cryosections and whole mount immunohistochemistry and imaging. CAL performed recombinant protein preparation, iPSC and vascularized organoid culture, generation of GBM biomimetics and TBI model, RNA-Seq, and manuscript writing. All authors contributed to manuscript preparation.

## REFERENCES

1 Lancaster, M. A. et al. Cerebral organoids model human brain development and microcephaly. Nature 501, 373–379, doi:10.1038/nature12517 (2013).

2 Lancaster, M. A. & Knoblich, J. A. Generation of cerebral organoids from human pluripotent stem cells. Nat Protoc 9, 2329–2340, doi:10.1038/nprot.2014.158 (2014).

3 Bagley, J. A., Reumann, D., Bian, S., Levi-Strauss, J. & Knoblich, J. A. Fused cerebral organoids model interactions between brain regions. Nat Methods 14, 743–751, doi:10.1038/nmeth.4304 (2017).

4 Sun, X. Y. et al. Generation of vascularized brain organoids to study neurovascular interactions. Elife 11, doi:10.7554/eLife.76707 (2022).

5 Cakir, B. et al. Expression of the transcription factor PU.1 induces the generation of microglia-like cells in human cortical organoids. Nat Commun 13, 430, doi:10.1038/s41467-022-28043-y (2022).

6 Lin, H. et al. A Scalable and Efficient Bioprocess for Manufacturing Human Pluripotent Stem Cell-Derived Endothelial Cells. Stem Cell Reports 11, 454–469, doi:10.1016/j.stemcr.2018.07.001 (2018).

7 Rhodes, C. T. et al. Region specific knock-out reveals distinct roles of chromatin modifiers in adult neurogenic niches. Cell Cycle, 1–13, doi:10.1080/15384101.2018.1426417 (2018).

8 Chang, K. C. et al. The chromatin repressors EZH2 and Suv4-20h coregulate cell fate specification during hippocampal development. FEBS Lett 596, 294–308, doi:10.1002/1873-3468.14254 (2022).

9 Whitesides, G. M. The origins and the future of microfluidics. Nature 442, 368–373, doi:10.1038/nature05058 (2006).

10 Klein, A. M. et al. Droplet barcoding for single-cell transcriptomics applied to embryonic stem cells. Cell 161, 1187–1201, doi:10.1016/j.cell.2015.04.044 (2015).

11 Taskesen, E. & Reinders, M. J. 2D Representation of Transcriptomes by t-SNE Exposes Relatedness between Human Tissues. PLoS One 11, e0149853, doi:10.1371/journal.pone.0149853 (2016).

12 Becht, E. et al. Dimensionality reduction for visualizing single-cell data using UMAP. Nat Biotechnol, doi:10.1038/nbt.4314 (2018).

13 Pollen, A. A. et al. Low-coverage single-cell mRNA sequencing reveals cellular heterogeneity and activated signaling pathways in developing cerebral cortex. Nat Biotechnol 32, 1053–1058, doi:10.1038/nbt.2967 (2014).

14 McKee, A. C. et al. TDP-43 proteinopathy and motor neuron disease in chronic traumatic encephalopathy. J Neuropathol Exp Neurol 69, 918–929, doi:10.1097/NEN.0b013e3181ee7d85 (2010).

15 McKee, A. C. et al. The spectrum of disease in chronic traumatic encephalopathy. Brain 136, 43–64, doi:10.1093/brain/aws307 (2013).

16 Goldstein, L. E. et al. Chronic traumatic encephalopathy in blast-exposed military veterans and a blast neurotrauma mouse model. Sci Transl Med 4, 134ra160, doi:10.1126/scitranslmed.3003716 (2012).

17 Rubenstein, R. et al. Tau phosphorylation induced by severe closed head traumatic brain injury is linked to the cellular prion protein. Acta Neuropathol Commun 5, 30, doi:10.1186/s40478-017-0435-7 (2017).

18 Rubenstein, R. et al. Comparing Plasma Phospho Tau, Total Tau, and Phospho Tau-Total Tau Ratio as Acute and Chronic Traumatic Brain Injury Biomarkers. JAMA Neurol 74, 1063–1072, doi:10.1001/jamaneurol.2017.0655 (2017).

19 Blennow, K. et al. Traumatic brain injuries. Nat Rev Dis Primers 2, 16084, doi:10.1038/nrdp.2016.84 (2016).

20 Shahim, P. et al. Serum neurofilament light protein predicts clinical outcome in traumatic brain injury. Sci Rep 6, 36791, doi:10.1038/srep36791 (2016).

21 Zetterberg, H. & Blennow, K. Fluid biomarkers for mild traumatic brain injury and related conditions. Nat Rev Neurol 12, 563–574, doi:10.1038/nrneurol.2016.127 (2016).

22 Lai, J. D. et al. KCNJ2 inhibition mitigates mechanical injury in a human brain organoid model of traumatic brain injury. Cell Stem Cell 31, 519–536 e518, doi:10.1016/j.stem.2024.03.004 (2024).

